# Investigating the effects of incremental conditioning and supplemental dietary tryptophan on the voluntary activity and behaviour of mid-distance training sled dogs

**DOI:** 10.1101/2020.04.21.052860

**Authors:** Eve Robinson, James R. Templeman, Emma Thornton, Candace C. Croney, Lee Niel, Anna K. Shoveller

## Abstract

Serotonin is a neurotransmitter synthesized by the amino acid tryptophan (Trp) that has the potential to impact the behaviour and activity of dogs. The objective of this study was to assess the effects of supplemental Trp and a 12-week incremental training regimen on the voluntary activity and behaviour of client-owned Siberian Huskies. Sixteen dogs were blocked for age, BW and sex and then randomly allocated to either the control or treatment group. Both groups were fed the same dry extruded diet; however, the treatment group were supplemented with Trp to achieve a Trp: large neutral amino acid ratio of 0.075:1. Once a week, a 5-minute video recording was taken immediately pre- and post- exercise to evaluate dogs’ behaviours. Activity monitors were used to record voluntary activity on both training and rest days. Linear regression analysis was used to assess the relationship between training week and time spent performing each behaviour. Additionally, a repeated measure mixed model was used to test differences between diet groups and training week for both behavioural and activity count data. The time spent performing agonistic behaviours prior to exercise was negatively associated with week for treatment dogs (β = −0.32, 95% CI [−0.55, −0.10], P < 0.05) and no change was observed for control dogs (β = −0.13, 95% CI [−0.41, 0.15], P > 0.10). Treatment did not have any effect on activity levels (P > 0.10). For all dogs, locomotive behaviours decreased prior to exercise as weeks progressed (P < 0.05), while run day voluntary activity depended on the distance run that day (P < 0.05). These data suggest that sled dogs experience an exercise-induced reduction in voluntary locomotion in response to both single bouts and repetitive bouts of exercise. Additionally, tryptophan supplementation may decrease agonistic behaviours, without having any effect on voluntary activity.

## Introduction

Sled dogs are endurance athletes that perform high levels of repetitive aerobic and resistance exercise. The success of sled dogs depends on multiple factors, including aerobic capacity and physical fitness as well as workability and trainability. While physical fitness is of primary importance, in order to achieve and maintain the level of fitness required to compete at a high level, it is critical to ensure that sled dogs continue to be motivated to exercise throughout their training and racing seasons. However, repetitive training programs can induce oxidative stress and muscle damage in working dogs [1, 2], which can result in muscle fatigue and soreness [3]. While researchers have examined the physiological effects of exercise and anticipation of exercise in sled dogs [4], more focus is needed on understanding how exercise can impact behaviour in both the short and long term. Exercise-induced muscle fatigue, as well as overtraining, can lead to decreased mood and lack of motivation to exercise in humans [5]. In canines, these symptoms may manifest as behavioural changes, such as a reduction in voluntary locomotive behaviours during an anticipatory period prior to a bout of exercise or a generalized decrease in voluntary daily activity. However, there is a dearth of literature that defines how commonly implemented training regimens and repetitive bouts of exercise may impact the behaviour of performance dogs.

Exercise capacity and motivation are not the only important factors to consider when investigating the performance of sled dogs. Sled dogs work in teams of 2 to 18 and typically interact with one or more handlers throughout their lives. Therefore, another key component to a successful sled dog is the ability to work in close proximity with other dogs and humans. Undesirable behavioural traits commonly reported in working dogs include fear, anxiety and both inter-dog and human-directed aggression [6, 7]. For sled dogs, aggression can present itself as contact or non-contact social conflict, which are forms of agonistic behaviours. These interactions can result in a poor team environment and may lead to pain and injury [7]. Previous research has found that dogs who display aggressive behaviours have lower serum and central serotonin concentrations than non-aggressive dogs [8, 9].

Serotonin, a neurotransmitter associated with regulation of mood, is synthesized in the brain from the amino acid tryptophan (Trp) [10]. Increased serotonin can enhance stress resistance [11] and reduce the prevalence of undesirable behaviours, such as anxiety [12], fear [13] and agonistic behaviours [14] in numerous monogastric species, including dogs [15]. Serotonergic activity is also linked to alterations in general voluntary activity and locomotion [16, 17]. Tryptophan has sedative effects in humans [18] and increasing central serotonin during exercise has been proposed to be associated with feelings of lethargy and a lack of motivation [19]. However, other research has presented conflicting results, with some reports indicating that increased levels of dietary Trp decreases fatigue perception in humans [20, 21], and has no effect on hyperactivity in client-owned dogs [15]. Increasing central serotonin concentrations may reduce various locomotory behaviours and activity; however, no previous research has looked at the effects of Trp-supplementation on the locomotive behaviours or voluntary activity levels in actively-training sled dogs.

Supplementing Trp in canine diets has been previously investigated as a means of increasing the production of serotonin [22, 15]. However, Trp competes with the large neutral amino acids (LNAA) for transport across the blood-brain barrier. Most protein-containing ingredients have lower concentrations of Trp relative to other amino acids, thus resulting in a reduced Trp:LNAA ratio [10]. Therefore, diet formulation, while still meeting the requirements for Trp, may lead to an imbalance of Trp and LNAA. Thus, the ratio of Trp:LNAA should be considered when formulating diets to ensure adequate transport of Trp to the brain for serotonin synthesis. The suggested Trp:LNAA ratio is 0.061:1 [23], although the optimal ratio may be greater [24] suggesting that current sporting dog diets may not adequately supply Trp to support serotonin synthesis. Therefore, increasing dietary Trp and the ratio of Trp:LNAA may result in an increase in central serotonin production and a reduction in agonistic or other undesirable behaviours in actively training sled dogs.

The objective of this study was to investigate the effects of 12 weeks of incremental conditioning and supplemental dietary Trp on the voluntary activity and pre and post- exercise gangline behaviours of mid-distance training sled dogs. We hypothesized that dietary supplementation of Trp would decrease the time spent performing various locomotive behaviours and agonistic behaviours, pre-and post-exercise, as well as decrease general voluntary activity, due to an increase in serotonergic activity. We also hypothesized that as exercise intensity and duration increase, the voluntary activity and locomotive behaviours performed would decrease due to exercise-induced physical fatigue.

## Materials and methods

### Animals, training regimen and diet

All procedures and facilities were approved by the Animal Care Committee at the University of Guelph (AUP #4008). Sixteen client-owned domestic Siberian Huskies (9 female: 4 intact, 5 spayed; 7 males: 2 intact, 5 neutered), with an average age of 4.8 ± 2.5 years and body weight (BW) of 24.3 ± 4.3kg, were housed, fed and trained at an off-site facility (RaJenn Siberian Huskies, Ayr, Ontario, Canada). A training regimen was proposed where dogs ran in a standard 16-dog gangline formation four times a week (Mon-Thurs) and distance increased incrementally over a 12-week period. Dogs ran in the same position on the gangline throughout the study. Dogs were anticipated to run 8km during week 0 and reach 86km during week 11; however, due to inclement weather, the training regimen was adjusted (Table 1; refer to Templeman et al. [25] for full proposed and adjusted training regimen). Daily temperature was recorded (Table 1). Total distance travelled over the training period was reduced from ~1900km to ~1230km. When training, dogs pulled an all-terrain vehicle carrying one passenger while maintaining an average speed of approximately 15km per hour throughout the study period. Training began consistently at 08:30 h. When not running, dogs were group housed in free-run outdoor kennels ranging from 3.5 to 80 square meters, containing anywhere from 2 – 10 dogs each. Two dogs were removed from the trial (one CON dog on week 7 and one TRT dog on week 9) due to exercise-related injuries. All data collected up until their respective points of removal are included in this report.

**Table 1.** Distance (km) run and ambient daily temperature (°C) when behavioural evaluations were carried out during 12 weeks of incremental conditioning for dogs fed either a treatment diet containing supplemental Trp compared to dogs fed control diet.

| Week | Distance (km) | Ambient temperature (°C) |
| --- | --- | --- |
| 0 | 8.9 | 5 |
| 1 | 12.9 | 7 |
| 2 | 19.7 | 3 |
| 3 | 26.7 | 7 |
| 4 | 30.8 | 3 |
| 5 | 38.4 | -2 |
| 6 | 30.2 | 3 |
| 7 | 30.0 | 3 |
| 8 | 30.0 | 1 |
| 9 | 53.2 | -5 |
| 10 | 30.2 | 0 |
| 11 | 38.2 | -4 |

Dogs were blocked for age, gender and BW and then randomly assigned into one of two groups (n = 8 per group): the control group (CON), fed a dry extruded diet (Champion Petfoods LT., Morinville, AB; refer to Templeman et al. [25] for full diet formulation) formulated to meet or exceed all AAFCO (2016) nutrient recommendations, or the treatment group (TRT), fed the control diet top-dressed with dietary Trp so as to reach a Trp:LNAA ratio of 0.075:1. Tryptophan solution was prepared by dissolving 10g of crystalline Trp (ADM Animal Nutrition, Woodstock, ON) per L of deionized water heated to 30°C. The solution was brought to room temperature (22 °C) and stored at 4°C until use. Each diet was stirred for 10 minutes after the addition of Trp solution to equally coat all kibble and ensure homogenous incorporation. Dogs were individually fed once a day at approximately 16:00 h to maintain BW, with individual BW recorded weekly. Dogs were provided *ad libitum* access to clean water.

### Behavioral evaluation

Using a digital camera (Sony HDR-CX110 HD Handycam, Sony Corp., Tokyo, Japan), video recordings were taken on one day (either Day 1 or Day 2) of each week to evaluate changes in the dogs pre- and post-exercise behaviours, where the distances they ran depended on week (Table 1). Once all dogs were put into their harnesses and individually placed in their respective positions on the 16-dog gangline, they were recorded for 5 continuous minutes immediately prior to exercise. Upon return from the training bout, 5 minutes of continuous video was again recorded while the dogs remained on the gangline. Dogs had been previously acclimatized to remain on the gangline post-exercise. Immediately upon cessation of the video, dogs were removed from the harness, and returned to their respective pens. Ten out of the 16 dogs (5 CON dogs and 5 TRT dogs) were chosen to be recorded, based on gangline position and visibility, to identify the occurrence of the following behaviours: jumping, lunging, changing posture, chewing on the gangline, sitting, lying, standing, digging, and changing posture (Table 2). All behavioural analysis was completed by a single individual who was blind to treatment groups. The overall time spent performing each behaviour (seconds) was determined from the video.

**Table 2.** Description of behavioural parameters analyzed during 5 minutes of video taken immediately pre- and post exercise for dogs fed either a treatment diet containing supplemental Trp compared to dogs fed control diet.

| Behaviour | Description |
| --- | --- |
| Sitting | Positioned with rear end and two front paws in contact with the ground |
| Lying | Positioned with ventral or side body in contact with the ground |
| Standing | Upright position with all four paws in contact with the ground |
| Digging | Using two front paws to dig at the ground |
| Posture changes | Frequent changes in state of motion, repeatedly lifting paws, pacing (>3 s) |
| Jumping | Upward motion where all four paws leave the ground |
| Lunging | Upward and forward motion where front two paws leave the ground |
| Agonistic Behaviour | Behaviours associated with social conflict, including noncontact (baring teeth, snapping) and contact (biting, nosing, wrestling) |

\*Behaviours classified as locomotive behaviours

### Activity monitoring

Three-dimensional accelerometers (Fitbarks, Fitbark Inc., Kansas City, MO) were attached to the collars of each dog to record activity on weeks 0, 6, and 11 (Table 1). For each of those weeks, activity was evaluated continuously for 24 hours during a rest day (no training) and again on an active day (training). Data is expressed as an activity count, which represents physical activity and is generated by company algorithm. While activity was being recorded, any periods of human interference, such as feeding, owner interaction, or training, were noted and subsequently removed from the activity count data. This ultimately left 3 hours of uninterrupted data that represented the voluntary activity performed by the dogs in their kennels, which was used for further analysis. For rest days, 3 consecutive hours of data were used from 11:00 h to 14:00h, while for active days, total activity counts were combined from 1-h pre-run (7:00 h to 8:00 h), 1-h post-run (dependent on run finishing time) and 1-h post-feeding (18:15h to 19:15h). Activity data from 4 CON dogs was removed from the week 1 rest day due to unanticipated owner interaction.

### Statistical analysis

The time spent performing a behaviour was converted into percentage of time [(duration of behaviour/duration of recording) × 100]. The average length of a bout of agonistic behaviour was also calculated [sum of duration of bouts/number of bouts]. The relationship between training week and the percentage of time performing a behaviour was analyzed using PROC REG of SAS (v.9.4; SAS Institute Inc., Cary, NC). If both TRT and CON groups had similar significant regression slopes for a particular behaviour, data were pooled and reanalyzed using PROC REG of SAS (v.9.4; SAS Institute Inc., Cary, NC). Behavioural data were also analyzed using PROC GLIMMIX of SAS (v.9.4; SAS Institute Inc., Cary, NC), with dog as a random effect and week and treatment as fixed effects. Week*treatment interaction effects were analyzed but removed if insignificant. Week was additionally treated as a repeated measure. Means were separated using Fisher’s LSD. Results are reported as least square means (LSM) ± standard error (SE). For all models, residuals were tested for homogeneity and normality by using the Shapiro-Wilk test and plots. PROC CORR of SAS (v.9.4; SAS Institute Inc., Cary, NC) was used to assess the relationship between daily temperature (°C), behaviour and week. Significance was declared at P ≤ 0.05, and trends at 0.05 < P ≤ 0.10.

Activity counts during rest days and training days were analyzed separately but using the same statistical method. Data were analyzed using the PROC GLIMMIX of SAS (v.9.4; SAS Institute Inc., Cary, NC) with dog as a random effect and week and treatment as fixed effects. Week was treated as a repeated measure. Week*treatment interaction effects were also analyzed. Means were separated using the Tukey adjustment. Results are reported as LSM ± SE. Additionally, PROC REG of SAS (v.9.4; SAS Institute Inc., Cary, NC) was used to evaluate the relationship between distance of exercise bout and run day activity counts. PROC CORR of SAS (v.9.4; SAS Institute Inc., Cary, NC) was used to assess the relationship between daily temperature (°C) and activity. For all models, residuals were tested for homogeneity and normality by using the Shapiro-Wilk test and plots. Significance was declared at P ≤ 0.05, and trends at 0.05 < P ≤ 0.10.

## Results

### Behaviour

#### Pre-exercise gangline behaviour

There was a negative association between the duration of a bout of agonistic behaviours and week of training for dogs receiving Trp-supplementation (β = −0.32, 95% CI [−0.55, −0.09], R^2^ = 0.10, P = 0.007; **Fig 1B**); however, no association was observed for dogs receiving the control diet (β = −0.13, 95% CI [−0.41, 0.15], R^2^ = 0.01, P > 0.10; **Fig 1A**). For all other behaviours, similar regression slopes were found between CON and TRT dogs; therefore, data were pooled to assess the effects of exercise. When the data were pooled, there was a negative association between week of training and time spent lunging and changing posture, and a positive association between week of training and time spent lying down (P < 0.05; **Table 3**). No associations were found between training week and time spent chewing on the line, digging, sitting or standing (P > 0.10; **Table 3**).

**Fig 1.**
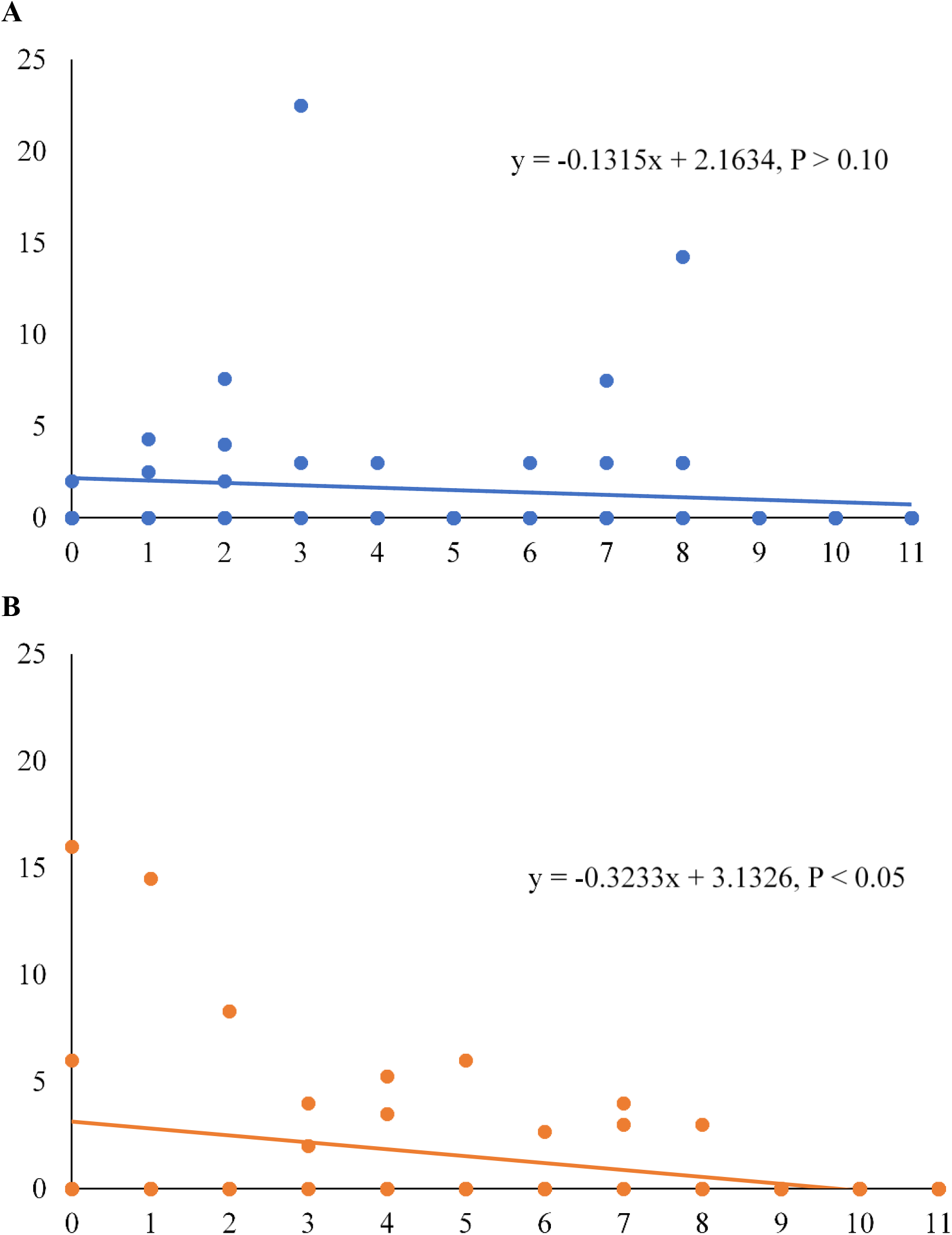
Agonistic behaviour performed by sled dogs undergoing 12 weeks of incremental conditioning. Average bout (sec) of agonistic behaviour performed by control dogs (A) and tryptophan-supplemented dogs (B) during 5-minutes immediately prior to exercise.

**Table 3.**
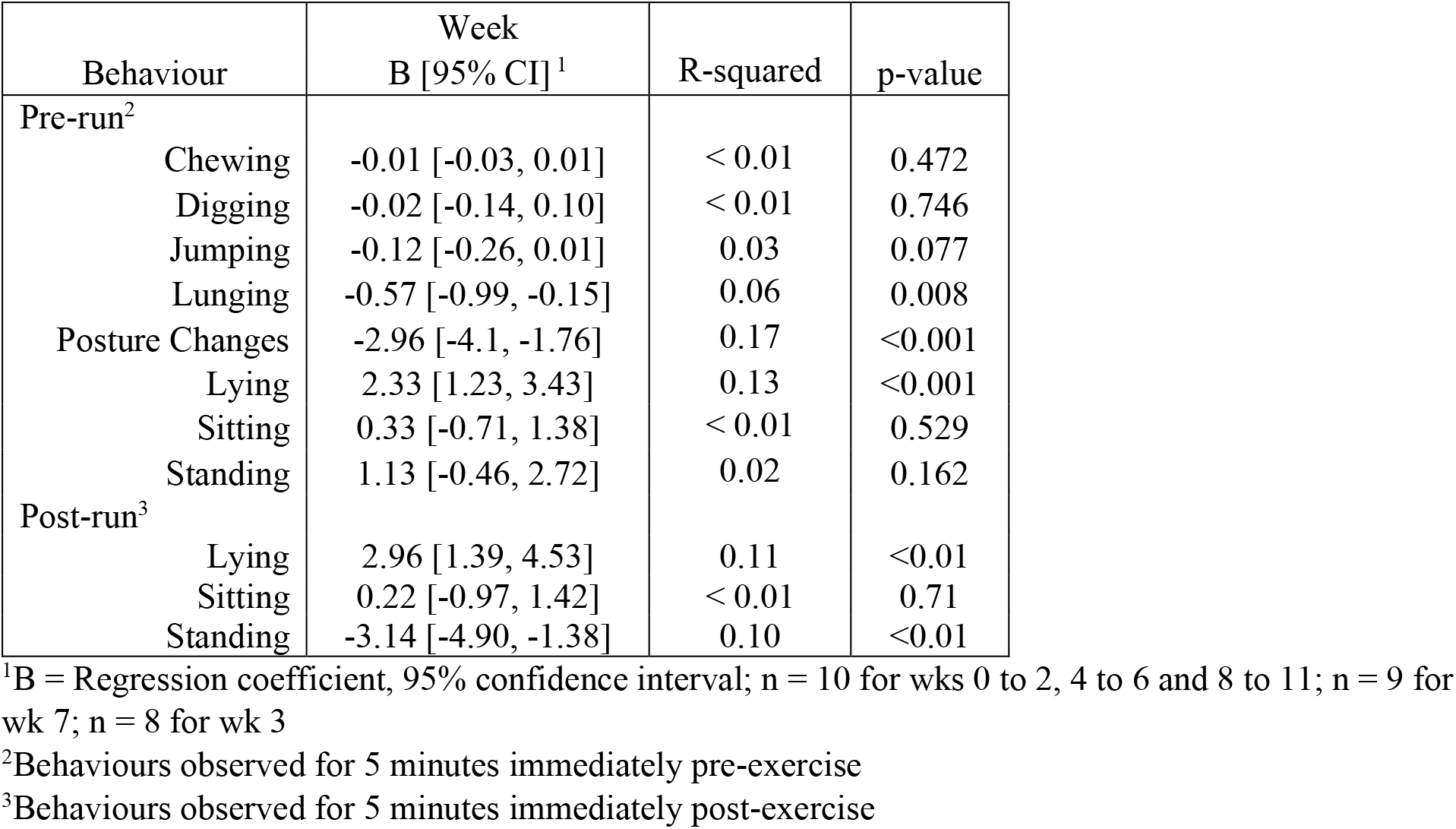
Linear regression estimates for the relationship between week of training and the time spent performing a behaviour for sled dogs undergoing 12 weeks of incremental conditioning.

| Behaviour | Week<br>B [95% CI]<sup>1</sup> | R-squared | p-value |
| --- | --- | --- | --- |
| Pre-run<sup>2</sup> |  |  |  |
| Chewing | -0.01 [-0.03, 0.01] | < 0.01 | 0.472 |
| Digging | -0.02 [-0.14, 0.10] | < 0.01 | 0.746 |
| Jumping | -0.12 [-0.26, 0.01] | 0.03 | 0.077 |
| Lunging | -0.57 [-0.99, -0.15] | 0.06 | 0.008 |
| Posture Changes | -2.96 [-4.1, -1.76] | 0.17 | <0.001 |
| Lying | 2.33 [1.23, 3.43] | 0.13 | <0.001 |
| Sitting | 0.33 [-0.71, 1.38] | < 0.01 | 0.529 |
| Standing | 1.13 [-0.46, 2.72] | 0.02 | 0.162 |
| Post-run<sup>3</sup> |  |  |  |
| Lying | 2.96 [1.39, 4.53] | 0.11 | <0.01 |
| Sitting | 0.22 [-0.97, 1.42] | < 0.01 | 0.71 |
| Standing | -3.14 [-4.90, -1.38] | 0.10 | <0.01 |

<sup>1</sup>B = Regression coefficient, 95% confidence interval; n = 10 for wks 0 to 2, 4 to 6 and 8 to 11; n = 9 for wk 7; n = 8 for wk 3
<sup>2</sup>Behaviours observed for 5 minutes immediately pre-exercise
<sup>3</sup>Behaviours observed for 5 minutes immediately post-exercise

There was no effect of dietary treatment on the overall time spent performing any behaviour evaluated (P > 0.10; S1 Table), therefore data from all dogs were pooled to examine week by week differences in behaviour. The time spent lunging was greater during week 1 than weeks 0, 6, 7, 9, 10 and 11, greater during week 2 than weeks 7, 9 and 11, and greater during weeks 3 and 4 than weeks 7 and 9 (P = 0.025; **Table 4**). Time spent changing posture was greater during weeks 0, 3, 4, 5 and 6 than weeks 9 to 11, and greater during weeks 1 and 2 than weeks 5 to 11 (P < 0.001; **Table 4**). Time spent lying down was greater during week 9 than any other week, and greater during weeks 7 and 11 than weeks 0 to 2 and 4 to 6 (P < 0.001; **Table 4**). Week tended to have an effect on time spent sitting (P = 0.065; **Table 4**) but had no effect on time spent performing agonistic behaviours, chewing on the gangline, digging, jumping or standing (P > 0.10; **Table 4**).

**Table 4.** Average percent of time (%) spent performing observed behaviours during 5-min pre exercise throughout 12 weeks of incremental conditioning.

| Behaviour | Week (Distance ran) |  |  |  |  |  |  |  |  |  |  |  | SEM<sup>1</sup> | p-value |
| --- | --- | --- | --- | --- | --- | --- | --- | --- | --- | --- | --- | --- | --- | --- |
|  | 0<br>(8.9<br>km) | 1<br>(12.9<br>km) | 2<br>(19.7<br>km) | 3<br>(26.7<br>km) | 4<br>(30.8<br>km) | 5<br>(38.4<br>km) | 6<br>(30.2<br>km) | 7<br>(30<br>km) | 8<br>(30<br>km) | 9<br>(53.2<br>km) | 10<br>(30.2<br>km) | 11<br>(38.2<br>km) |  |  |
| Agonistic | 0.8 | 1.6 | 1.8 | 2.4 | 1.0 | 0.2 | 0.9 | 0.5 | 2.2 | 0 | 0 | 0 | 1.0 | 0.542 |
| Chewing | 0 | 0.1 | 0.1 | 0 | 0 | 0.4 | 0 | 0 | 0 | 0 | 0 | 0 | 0.1 | 0.318 |
| Digging | 0.1 | 0.6 | 0.4 | 0.7 | 1.3 | 0.1 | 2.3 | 0 | 0.2 | 0 | 0.8 | 0 | 0.8 | 0.551 |
| Jumping | 1 | 1.7 | 1.9 | 1.5 | 1.7 | 1.1 | 1 | 0 | 1.8 | 0 | 0.5 | 0.3 | 0.9 | 0.194 |
| Lunging | 2.5<sup>bcd</sup> | 9<sup>a</sup> | 7.6<sup>ab</sup> | 7.0<sup>abc</sup> | 6.0<sup>abc</sup> | 3.3<sup>abcd</sup> | 2.8<sup>bcd</sup> | 0<sup>d</sup> | 4.5<sup>abcd</sup> | 0<sup>d</sup> | 2.0<sup>bcd</sup> | 1.0<sup>cd</sup> | 2.8 | 0.025 |
| Postural<br>Changes | 32.0<sup>ab</sup> | 44.2<sup>a</sup> | 41.7<sup>a</sup> | 32.7<sup>ab</sup> | 34.7<sup>ab</sup> | 32.5<sup>ab</sup> | 25.2<sup>bc</sup> | 22.4<sup>bcd</sup> | 25.7<sup>bc</sup> | 2.8<sup>e</sup> | 13.7<sup>cde</sup> | 12.3<sup>de</sup> | 7.6 | <0.001 |
| Sitting | 22.5 | 1.7 | 10.8 | 6.3 | 1.7 | 1.6 | 5.5 | 2.5 | 5.1 | 20.7 | 17.2 | 11.8 | 6.8 | 0.065 |
| Standing | 39.3 | 39.6 | 36.1 | 44.4 | 51.1 | 59.9 | 63.3 | 55.0 | 56.6 | 29.5 | 53.1 | 55.5 | 10.4 | 0.134 |
| Lying | 1.9<sup>c</sup> | 1.8<sup>c</sup> | 0<sup>c</sup> | 4.2<sup>bc</sup> | 2.4<sup>c</sup> | 0.8<sup>c</sup> | 0<sup>c</sup> | 19.3<sup>b</sup> | 3.8<sup>bc</sup> | 47<sup>a</sup> | 12.7<sup>bc</sup> | 19.0<sup>b</sup> | 6.6 | <0.001 |

<sup>1</sup>Standard error of the mean; n = 10 for wks 0 to 2, 4 to 6 and 8 to 11; n = 9 for wk 7; n = 8 for wk 3
<sup>abc</sup>Values in a row with a different superscript are different (P < 0.05)

Environmental temperature was positively correlated with the time spent performing agonistic behaviours (r = 0.22, P = 0.0178), lunging (r = 0.27, P = 0.003), changing posture (r = 0.38, P <0.001) and negatively correlate with the time spent lying down (r = −0.41, P <0.001). Environmental temperature tended to be positively associated with time spent jumping (r = 0.17, P = 0.066), and was not correlated with time spent chewing, digging, sitting or standing (P > 0.10). Environmental temperature was negatively correlated with week (r = −0.75, P <0.001).

#### Post-exercise gangline behaviour

The only behaviours observed post-exercise were sitting, standing or lying. Similar regression slopes were found between CON and TRT dogs for the time spent performing any behaviour and week; therefore, data from all dogs were pooled. There was a positive association between week of training and time spent lying down and a negative association was found between week of training and time spent standing (P < 0.05; **Table 3**). No association was found between week and time spent sitting (P > 0.10; **Table 3**).

There was no effect of dietary treatment on the overall time spent performing any behaviour evaluated (P > 0.10; S2 Table), therefore data from all dogs were pooled to examine week by week differences in behaviour. The time spent standing was greater during weeks 0, 1, 2, 4 and 7 than weeks 8 and 11, greater during week 3 than weeks 8 to 11, greater during week 5 than week 8 and 11 and greater during week 6 than weeks 5 and 8 to 11 (P < 0.05; **Table 5**). Time spent lying down was greater during weeks 8 and 11 than weeks 0 - 7 and 9 (P < 0.05; **Table 5**). Week had no effect on time spent sitting (P > 0.10; **Table 5**).

**Table 5.** Average percent of time (%) spent performing observed behaviours during 5-min post exercise throughout 12 weeks of incremental conditioning.

| Behaviour | Week (distance ran) |  |  |  |  |  |  |  |  |  |  |  | SEM<sup>1</sup> | p-value |
| --- | --- | --- | --- | --- | --- | --- | --- | --- | --- | --- | --- | --- | --- | --- |
|  | 0<br>(8.9<br>km) | 1<br>(12.9<br>km) | 2<br>(19.7<br>km) | 3<br>(26.7<br>km) | 4<br>(30.8<br>km) | 5<br>(38.4<br>km) | 6<br>(30.2<br>km) | 7<br>(30<br>km) | 8<br>(30<br>km) | 9<br>(53.2<br>km) | 10<br>(30.2<br>km) | 11<br>(38.2<br>km) |  |  |
| Sitting | 11.8 | 10.8 | 14.5 | 10.2 | 4.1 | 13.9 | 6.7 | 4.1 | 13.0 | 20.1 | 12.8 | 12.1 | 8.2 | 0.83 |
| Standing | 75.3<sup>abc</sup> | 74.5<sup>abc</sup> | 70.8<sup>abc</sup> | 88.1<sup>ab</sup> | 70.0<sup>abc</sup> | 64.0<sup>bc</sup> | 91.0<sup>a</sup> | 75.9<sup>abc</sup> | 35.3<sup>d</sup> | 57.8<sup>cd</sup> | 55.4<sup>cd</sup> | 35.9<sup>d</sup> | 11.2 | <0.01 |
| Lying | 12.9<sup>bcd</sup> | 15.2<sup>bcd</sup> | 14.6<sup>bcd</sup> | 2.4<sup>cd</sup> | 24.9<sup>bcd</sup> | 25.6<sup>bc</sup> | 2.3<sup>d</sup> | 19.8<sup>bcd</sup> | 51.7<sup>a</sup> | 24.5<sup>bcd</sup> | 31.8<sup>ab</sup> | 52.0<sup>a</sup> | 10.0 | <0.01 |

<sup>1</sup>Standard error of the mean; n = 10 for wks 0 to 2, 4 to 6 and 8 to 11; n = 9 for wk 7; n = 8 for wk 3
<sup>abc</sup>Values in a row with a different superscript are different (P < 0.05)

Environmental temperature was negatively correlated with time spent lying (r = −0.30, P < 0.05), and positively associated with time spent standing (r = 0.31, P <0.05). Environmental temperature was not correlated with time spent sitting (r = −0.09, P > 0.10).

#### Activity

Treatment had no effect on activity levels during rest days or active days throughout the 12-week conditioning period (P > 0.10; **Table 6**). When data from all dogs were pooled, total activity levels on rest days (no regimented exercise) decreased from week 0 to week 6 and from week 6 to week 11 (P < 0.05; **Fig 2A**). Total activity levels on run days (regimented exercise) decreased between week 0 and 6 (P < 0.05); however, run day activity levels on week 11 did not differ from either week 0 or 6 (P > 0.05; **Fig 2B**). Additionally, total activity on active days was negatively associated with the distance run that day (β = −14.59, 95% CI [−22.03, −7.15], P < 0.05).

**Table 6.** Mean voluntary activity counts for control dogs or tryptophan-supplemented (treatment) dogs on active days and rest days during weeks 0, 6 and 11 of a 12-week incremental conditioning period.

|  | Week (Distance ran) |  |  | p-value |  |  |  |
| --- | --- | --- | --- | --- | --- | --- | --- |
|  | 0<br>(8.9 km) | 6<br>(46.2 km) | 11<br>(38.2 km) | SEM<sup>1</sup> | Week | Treatment | Week*Treatment |
| Active Day |  |  |  |  |  |  |  |
| Control | 1054.36<sup>a</sup> | 409.08<sup>ab</sup> | 538.34<sup>b</sup> | 165.52 | 0.03 | 0.73 | 0.51 |
| Treatment | 849.75<sup>a</sup> | 336.69<sup>b</sup> | 659.58<sup>ab</sup> | 148.11 | 0.03 |  |  |
| Rest Day |  |  |  |  |  |  |  |
| Control | 1358.02<sup>a</sup> | 1076.18<sup>a</sup> | 374.40<sup>b</sup> | 197.63 | <0.01 | 0.78 | 0.28 |
| Treatment | 1539.00<sup>a</sup> | 716.93<sup>b</sup> | 345.43<sup>b</sup> | 230.80 | <0.01 |  |  |

<sup>1</sup>Standard error of the mean; For active days, n = 8 for control and treatment week 0 and 6, n = 7 for control and treatment week 11; for rest days, n = 4 for control week 0, n = 8 for treatment week 0, n = 8 for control and treatment week 6, n = 7 for control and treatment week 11
<sup>abc</sup>Values in a row with a different superscript are different (P < 0.05)

**Fig 2.**
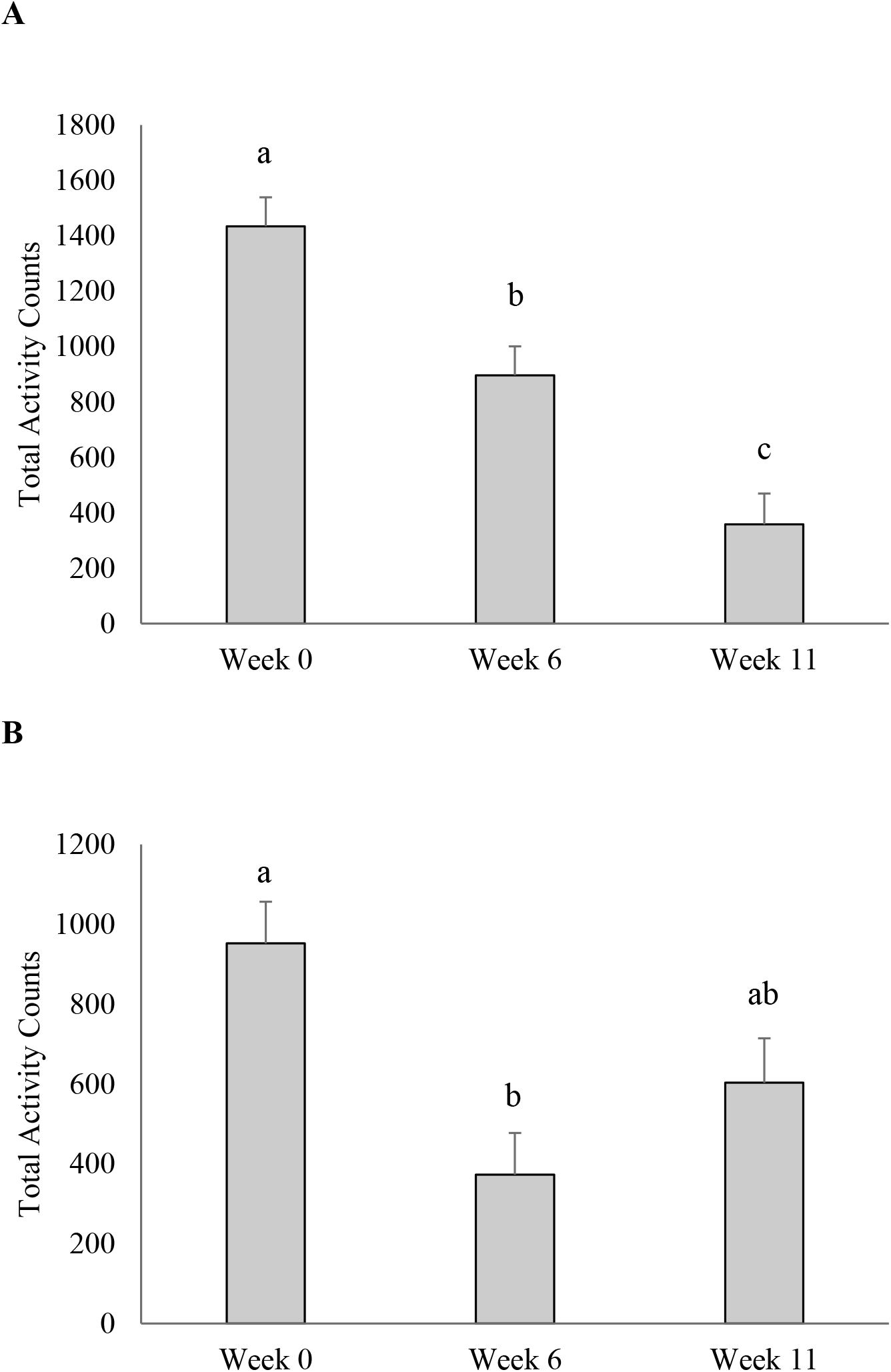
Average activity counts of sled dogs undergoing 12 weeks of incremental conditioning. (A) Average activity counts of sled during rest day (no regimented exercise). (B) Average activity counts during active days (regimented exercise) where dogs ran 8.9km, 46.2km and 38.2km during weeks 0, 6 and 11, respectively. Columns with different letters are different from each other (P < 0.05). Error bars represent the standard error of the mean.

Daily environmental temperature was positively correlated with off day activity (r = 0.62, P < 0.05) and run day activity (r = 0.31, P < 0.05).

## Discussion

To the authors’ knowledge, this is the first study to evaluate the effects of dietary Trp and an incremental training regimen on the pre- and post-exercise behaviour and voluntary activity of dogs. Dietary Trp may influence behaviours related to anxiety [12], stress [11] and fear [13] and in the present study, dogs receiving Trp supplementation experienced a reduction in agonistic behaviours prior to exercise throughout a 12-week period. However, Trp-supplementation did not affect any other observed behaviour, nor did it affect the voluntary activity of dogs during training or rest days. Although previous research has suggested that dietary Trp can have a sedative effect resulting in a decrease in locomotion in humans [18] and mice [17], we found no effect in actively training dogs. In agreement with the present work, research by DeNapoli et al. [15] found that dietary Trp supplementation did not reduce hyperactivity in dogs; which the authors characterized by criteria including excessive pacing, chewing of objects and the inability to remain in a sit position. These behaviours are comparable to the pre-exercise behaviours observed in the current study, which were also unaffected by dietary Trp supplementation. Additionally, Bosch et al. [22] found no differences in the percent of time spent changing locomotive states, walking or lying down in an open-field test following 8 weeks of Trp supplementation in client-owned dogs. Combined, these results suggest that Trp supplementation does not influence activity levels and associated locomotive behaviours in domestic dogs.

### Tryptophan effect on aggression-related behaviours

Although Trp did not appear to affect overall locomotion in sled dogs, there was a significant effect of the dietary treatment on agonistic behaviours performed throughout the study period. A decrease in aggression following dietary Trp supplementation has been previously reported in pigs [26] chickens [27], vervet monkeys [28] and rats [14]. As well, dogs previously diagnosed with aggression that were being fed a high protein diet supplemented with additional Trp to reach a Trp:LNAA of 0.07:1 showed reduced aggressive behaviour [15]. In the present study, no preliminary evaluations were performed to identify any potential behavioural pre-dispositions; however, Trp-supplementation still decreased the time spent performing agonistic behaviours prior to exercise over the 12-week period. The Trp:LNAA ratio of 0.075:1 in the treatment diet resulted in increased serum Trp concentration in the treatment dogs compared to the control dogs [25], which favors the conditions for higher transport of Trp across the blood-brain barrier. Theoretically, this suggests that more Trp will be converted to serotonin in the brain which is thought to be involved in the regulation of aggressive behaviour, likely through the enhancement of impulse control [29]. Male rhesus macaques with low levels of cerebrospinal fluid 5-hydroxyindoleacetic acid (CSF 5-HIAA), which is the primary metabolite of serotonin, were found to be more likely to show aggressive behaviours as well as experience a loss of impulse control characterized by greater risk-taking behaviours [30]. Dogs who demonstrated impulse aggression, identified by biting without warning, also had lower concentrations of CSF 5-HIAA than non-aggressive dogs [31]. Through the action of central serotonin, treatment dogs in our study may have experienced improved impulse control causing the slight reduction in agonistic behaviours prior to exercise. While the recommended Trp:LNAA ratio is 0.061:1 [23], the ratio needed to influence a behavioural response may be higher. Various inclusion levels of Trp in canine diets leading to Trp:LNAA ratios of 0.0274:1, 0.0403:1, 0.0448:1, and 0.0581:1, all had no impact on aggressive behaviours in response to a familiar or unfamiliar human in mixed-breed hounds [32]. However, a Trp:LNAA ratio of 0.075:1 used in the present study, and 0.07:1 used by DeNapoli et al. [15] both caused reductions in agonistic behaviours. This suggests that diets aimed to reduce agonistic behaviours should be formulated to include a Trp:LNAA ratio of over 0.07:1 in order to elicit the desired behavioural change. The results of this study further contribute to the existing literature that dietary Trp can influence behaviours related to aggression, with no apparent sedative effects on locomotive activity or behaviour. In the present study, the average level of agonistic behaviour was minimal during any given week (< 5%). Future research should consider incorporating a baseline evaluation to ensure adequate inclusion of dogs that are known to exhibit agonistic behaviours.

### Effect of single bout and repetitive exercise on behaviour and activity

In addition to the effects of dietary treatment on agonistic behaviours, additional findings from this study revealed that a 12-week conditioning period influenced the locomotive behaviours and voluntary activity of sled dogs, regardless of dietary treatment. Endurance exercise causes physiological changes, and recovery from exercise involves restoration of endogenous energy stores and a return to a normal heart rate, respiratory rate, and internal temperature [33]. In addition to the inevitable physiological impact, intense aerobic exercise in human athletes can lead to an increased risk of oxidative stress and the subsequent skeletal muscle damage is associated with soreness, reduced range of motion, and muscle fatigue [3, 34]. These physiological effects and the extent of recovery depend on the duration and frequency of exercise [35, 36]. In the present study, the amount of voluntary activity performed by the dogs in their free-run kennels during days of active training depended upon the distance run that day, which likely represents a decrease in available metabolic energy due to exercise. During the post-exercise period, dogs are likely resting, in part, to restore endogenous energy and return to homeostasis. Furthermore, as indicators of oxidative stress and muscle damage are linked to the intensity of aerobic exercise [37, 38], it is expected that voluntary activity performed by sled dogs would be dependent on the duration of an exercise bout. This suggests that voluntary activity surrounding exercise may be useful as an additional indicator of intensity of a bout of exercise in sled dogs.

While voluntary activity was found to be related to the duration of a single bout of exercise, it is also evident that behaviour and locomotion may be additionally affected by repetitive exercise. For all dogs, there was an overall decrease in voluntary activity during rest days and a reduction in locomotive behaviours pre-exercise over the 12-week study period. Although no markers of metabolic stress were measured in the present study, previous research has shown that repetitive exercise in sled dogs is associated with increased creatine kinase (CK) concentrations, which is an indirect marker of skeletal muscle damage [39]. Sled dogs who ran 58-km on each of three consecutive days had a significant increase in serum CK following the first exercise bout, with a further significant increase after the third exercise bout [1]. Although the distances run in the present study did not exceed 53km a day, it is possible that similar muscle fatigue was experienced, which caused the documented reduction in voluntary activity and locomotive behaviours. Taken together, this suggests that while short-term voluntary activity is related to the distance of an exercise bout on a specific day, activity and locomotion can also be affected in the long term by the repetitiveness of a training regimen. Sled dogs are simultaneously experiencing both an acute and chronic response to exercise, which should be considered when designing training regimens to optimize exercise programs. It is important to note that ambient temperature may also affect behaviour. Interestingly, in the present study, sled dogs were less active and exhibited fewer locomotive behaviours as temperatures decreased. Since the average daily temperature decreased as the study progressed, it is possible that these behavioural changes represent the effects of the repetitive training regimen rather than the decrease in temperature.

Along with the physiological impacts of endurance exercise, dogs may have also experienced additional psychological symptoms throughout the conditioning period. Although acute exercise is associated with a positive effect on mood [40], the extreme exertion experienced by sled dogs may have variable effects. In humans, over-training is most often characterized by a decrease in mood, lack of motivation to exercise and chronic fatigue [5, 41]. Although these symptoms are difficult to evaluate in working canines, it is likely that sled dogs have the potential to experience similar effects of over-training. Unfortunately, no previous research has defined behavioural signs of motivation to exercise in dogs. It is possible that the prevalence of postural changes or lunging forward on the gangline could be an indicator of anticipation or motivation to exercise when exhibited prior to running. The decrease in these behaviours seen throughout the conditioning period could potentially indicate a decrease in motivation; however, more research is needed to determine how these behaviours are associated with other indicators of fatigue and well-being. It is also possible that a decrease in these behaviours could suggest that dogs were becoming familiarized with the training regimen, therefore were less responsive as it progressed. Tracking observable behaviours prior to exercise may be useful for mushers or other sporting dog owners as an indication of motivation to exercise.

## Conclusion

The findings of the current study suggest that Trp supplementation decreases agonistic behaviours in actively training sled dogs, while having no effect on activity or locomotion. Future research should continue to examine the use of Trp to decrease pre-run agonistic behaviours in working dogs, with a focus on dogs who have been pre-diagnosed with behavioural issues. Additionally, the reduction in activity and locomotive behaviours, such as lunging and changes in posture, following exercise was related to both to the intensity of a bout of exercise as well as to the repetitiveness of the training regimen. Short-term voluntary activity was related to the distance of the bout of exercise performed that day, while repetitive exercise caused a progressive decrease in locomotive behaviours during a pre- and post- exercise period.

Future research should focus on assessing correlations between behavioural responses to repetitive exercise and physiological markers of over-training or fatigue. Overall, this research is the first to show the positive impact of an increased Trp:LNAA ratio on agonistic behaviour, which ultimately improves the workability of sled dogs and potentially decreases their risk of pain and injury. These results can be used to inform the development of diets and training programs designed to maximize the performance and success of sled dogs.

## References

1. Hinchcliff KW, Reinhart GA, DiSilvestro R, Reynolds A, Blostein-Fujii A, Swenson RA. Oxidant stress in sled dogs subjected to repetitive endurance exercise. Am J Vet Res. 2000 May;61(5): 512–7.

2. Pasquini A, Luchetti E, Cardini G. Evaluation of oxidative stress in hunting dogs during exercise. Res Vet Sci. 2010 Aug 1;89(1):120–3.

3. Ji LL, Leichtweis S. Exercise and oxidative stress: Sources of free radicals and their impact on antioxidant systems. Age (Omaha). 1997 Apr;20(2):91–106.

4. Angle CT, Wakshlag JJ, Gillette RL, Stokol T, Geske S, Adkins TO, et al. Hematologic, serum biochemical, and cortisol changes associated with anticipation of exercise and short duration high-intensity exercise in sled dogs. Veterinary Clinical Pathology. 2009;38(3):370–4.

5. Stone MH, Keith RE, Kearney JT, Fleck SJ, Wilson GD, Triplett NT. Overtraining: A Review of the Signs, Symptoms and Possible Causes. The Journal of Strength & Conditioning Research. 1991 Feb;5(1):35.

6. Haverbeke A, De Smet A, Depiereux E, Giffroy J-M, Diederich C. Assessing undesired aggression in military working dogs. Applied Animal Behaviour Science. 2009 Feb 1;117(1):55–62.

7. Rooney NJ, Clark CCA, Casey RA. Minimizing fear and anxiety in working dogs: A review. Journal of Veterinary Behavior. 2016 Nov;16:53–64.

8. Çakiroǧlu D, Meral Y, Sancak AA, Çifti G. Relationship between the serum concentrations of serotonin and lipids and aggression in dogs. Veterinary Record. 2007 Jul 14;161(2):59–61.

9. León M, Rosado B, García-Belenguer S, Chacón G, Villegas A, Palacio J. Assessment of serotonin in serum, plasma, and platelets of aggressive dogs. Journal of Veterinary Behavior. 2012 Nov;7(6):348–52.

10. Leathwood PD. Tryptophan Availability and Serotonin Synthesis. Proceedings of the Nutrition Society. 1987 Feb;46(01):143–56.

11. Koopmans SJ, Ruis M, Dekker R, van Diepen H, Korte M, Mroz Z. Surplus dietary tryptophan reduces plasma cortisol and noradrenaline concentrations and enhances recovery after social stress in pigs. Physiology & Behavior. 2005 Jul 21;85(4):469–78.

12. Orosco M, Rouch C, Beslot F, Feurte S, Regnault A, Dauge V. Alpha-lactalbumin-enriched diets enhance serotonin release and induce anxiolytic and rewarding effects in the rat. Behavioural Brain Research. 2004 Jan;148(1-2):1–10.

13. Rouvinen K, Archbold S, Laffin S, Harri M. Long-term effects of tryptophan on behavioural response and growing-furring performance in silver fox (Vulpes vulpes). Applied Animal Behaviour Science. 1999 Mar 1;63(1):65–77.

14. Gibbons JL, Barr GA, Bridger WH, Leibowitz SF. L-tryptophan’s effects on mouse killing, feeding, drinking, locomotion, and brain serotonin. Pharmacology Biochemistry and Behavior. 1981 Aug;15(2):201–6.

15. DeNapoli JS, Dodman NH, Shuster L, Rand WM, Gross KL. Effect of dietary protein content and tryptophan supplementation on dominance aggression, territorial aggression, and hyperactivity in dogs. Journal of the American Veterinary Medical Association. 2000 Aug 1;217(4):504–8.

16. Gainetdinov RR, Wetsel WC, Jones, Levin ED, Jaber M, Caron MG. Role of serotonin in the paradoxical calming effect of psychostimulants on hyperactivity. Science (Washington). 1999;283(5400):397–401.

17. Lasley SM, Thurmond JB. Interaction of dietary tryptophan and social isolation on territorial aggression, motor activity, and neurochemistry in mice. Psychopharmacology. 1985 Nov;87(3):313–21.

18. Leathwood PD, Pollet P. Diet-induced mood changes in normal populations. J Psychiatr Res. 1982 1983;17(2):147–54.

19. Newsholme EA, Blomstrand E. Branched-Chain Amino Acids and Central Fatigue. J Nutr. 2006 Jan 1;136(1):274S–276S.

20. Segura R, Ventura JL. Effect of L-tryptophan supplementation on exercise performance. Int J Sports Med. 1988 Oct;9(5):301–5.

21. Javierre C, Segura R, Ventura JL, Suárez A, Rosés JM. L-Tryptophan Supplementation Can Decrease Fatigue Perception During an Aerobic Exercise with Supramaximal Intercalated Anaerobic Bouts in Young Healthy Men. International Journal of Neuroscience. 2010 Apr 1;120(5):319–27.

22. Bosch G, Beerda B, Beynen AC, van der Borg JAM, van der Poel AFB, Hendriks WH. Dietary tryptophan supplementation in privately owned mildly anxious dogs. Applied Animal Behaviour Science. 2009 Dec;121(3-4):197–205.

23. National Research Council. Nutrient Requirements for dogs and cat. 2^nd^rev. ed. Washington, DC: The National Academies Press; 2006.

24. Templeman J, Mansilla W, Fortener L, Shoveller A. 371 Tryptophan requirements in small, medium, and large breed adult dogs using the indicator amino acid oxidation technique. J Anim Sci. 2018 Dec 7;96(suppl_3):148–148.

25. Templeman J, Thornton E, Cargo-Froom C, Squires E, Swanson K, Shoveller, A. Effects of incremental exercise and dietary tryptophan supplementation on the amino acid metabolism, serotonin status, stool quality, fecal metabolites, and body composition of mid-distance trained sled dogs. J Anim Sci. 2020; Forthcoming.

26. Li YZ, Kerr BJ, Kidd MT, Gonyou HW. Use of supplementary tryptophan to modify the behavior of pigs. Journal of animal science. 2006;84(1):212–20.

27. Shea MM, Douglass LW, Mench JA. The interaction of dominance status and supplemental tryptophan on aggression in Gallus domesticus males. Pharmacology Biochemistry and Behavior. 1991 Mar 1;38(3):587–91.

28. Chamberlain B, Ervin FR, Pihl RO, Young SN. The effect of raising or lowering tryptophan levels on aggression in vervet monkeys. Pharmacology Biochemistry and Behavior. 1987 Dec 1;28(4):503–10.

29. Coccaro EF. Central serotonin and impulsive aggression. Br J Psychiatry Suppl. 1989 Dec;(8):52–62.

30. Mehlman PT, Higley JD, Faucher I, Lilly AA, Taub DM, Vickers J, et al. Low CSF 5-HIAA concentrations and severe aggression and impaired impulse control in nonhuman primates. The American Journal of Psychiatry. 1994;151(10):1485–91.

31. Reisner IR, Mann JJ, Stanley M, Huang Y, Houpt KA. Comparison of cerebrospinal fluid monoamine metabolite levels in dominant-aggressive and non-aggressive dogs. Brain Research. 1996 Apr;714(1-2):57–64.

32. Templeman JR, Davenport GM, Cant JP, Osborne VR, Shoveller A-K. The effect of graded concentrations of dietary tryptophan on canine behavior in response to the approach of a familiar or unfamiliar individual. Can J Vet Res. 2018 Oct;82(4):294–305.

33. Gillette RL, Angle TC, Sanders JS, DeGraves FJ. An evaluation of the physiological affects of anticipation, activity arousal and recovery in sprinting Greyhounds. Applied Animal Behaviour Science. 2011 Mar;130(3-4):101–6.

34. Peternelj T-T, Coombes JS. Antioxidant Supplementation during Exercise Training: Beneficial or Detrimental? Sports Medicine. 2011 Dec;41(12):1043–69.

35. Noakes TD. Effect of Exercise on Serum Enzyme Activities in Humans. Sports Medicine. 1987 Jul 1;4(4):245–67.

36. Nieman DC, Dumke CL, Henson DA, McAnulty SR, Gross SJ, Lind RH. Muscle damage is linked to cytokine changes following a 160-km race. Brain Behav Immun. 2005 Sep;19(5):398–403.

37. Skenderi KP, Kavouras SA, Anastasiou CA, Yiannakouris N, Matalas A-L. Exertional Rhabdomyolysis during a 246-km Continuous Running Race: Medicine & Science in Sports & Exercise. 2006 Jun;38(6):1054–7.

38. Ramos D, Martins EG, Viana-Gomes D, Casimiro-Lopes G, Salerno VP. Biomarkers of oxidative stress and tissue damage released by muscle and liver after a single bout of swimming exercise. Appl Physiol Nutr Metab. 2013 Apr 20;38(5):507–11.

39. Mckenzie E, Holbrook T, Williamson K, Royer C, Valberg S, Hinchcliff K, et al. Recovery of Muscle Glycogen Concentrations in Sled Dogs during Prolonged Exercise: Medicine & Science in Sports & Exercise. 2005 Aug;37(8):1307–12.

40. Menor-Campos DJ, Molleda-Carbonell JM, López-Rodríguez R. Effects of exercise and human contact on animal welfare in a dog shelter. Veterinary Record. 2011 Oct 8;169(15):388–388.

41. McKenzie DC. Markers of excessive exercise. Canadian Journal of Applied Physiology. 1999 Feb;24(1):66-.

